# Repeated low-intensity focused ultrasound induces microglial profile changes in the TgF344-AD rat model of Alzheimer’s disease

**DOI:** 10.1101/2024.09.25.614692

**Authors:** Laurene Abjean, Anthony Novell, Benoît Larrat, Boris Rafael Gueorguiev, Thomas Cailly, Christine Fossey, Frédéric Fabis, Rares Salomir, Stergios Tsartsalis, Benjamin B. Tournier, Philippe Millet, Kelly Ceyzériat

**Author notes:** Corresponding author: Kelly Ceyzériat, PhD, CIBM Center for Biomedical Imaging University of Geneva, Avenue de la Roseraie, 64 1205 Geneva, Switzerland. These authors equally contributed to this work (Philippe Millet, Kelly Ceyzériat).

## Abstract

Alzheimer’s disease (AD), the most common cause of dementia, represents one of the main clinical challenges of the century as the number of patients is predicted to triple by 2050. Despite the recent approval of three monoclonal antibodies targeting Amyloid β (Aβ) aggregates by the Food and Drug Administration (FDA), immunotherapies still face challenges due to the difficulty of antibodies crossing the blood-brain barrier (BBB). This necessitates administering large doses of drugs to achieve their therapeutic effects, which is associated with significant side effects. In this context, low-intensity focused ultrasound (LiFUS) appears as an innovative and non-invasive method which, in association with intravenous injection of microbubbles (MB), leads to a transient BBB opening. This innovative strategy has been extensively studied in different preclinical models and more recently in human clinical trials, particularly in the context of AD. LiFUS+MB increases the inflammatory response at short-term, but the time course of this response is not consistent between studies, certainly due to the discrepancy between LiFUS protocols used. Moreover, the impact at longer term is understudied and the mechanisms underlying this effect are still not well understood. In our study, we therefore used the TgF344-AD rat model of AD to investigate the effect of a single or multiple exposures to LiFUS+MB in a large volume of the brain on inflammatory response, tauopathy and amyloid load, at both early and advanced stages. The ultrasound attenuation through the skull was corrected to apply a peak negative acoustic pressure of 450 kPa in all treated animals. At an advanced disease stage, single LiFUS+MB exposure induces a slight astrocyte and microglial response 24 hours post-treatment whereas chronic LiFUS treatment is associated with a transient inflammatory response predominantly affecting microglial cells, which is no longer detectable 6 weeks post-treatment. At an early stage of pathology, LiFUS seems to induce microglial reprogramming, leading to the adaptation of gene expression related to key functions such as inflammatory response, mitochondrial and energetic metabolism. In our rat model and LiFUS+MB protocol conditions, a single LiFUS exposure reduced significantly highly aggregated Aβ42 peptide concentration. Surprisingly, multiple exposures had this opposite effect at short-term but not at longer term.

## Introduction

Alzheimer’s disease (AD), one of the main causes of dementia, is associated at the neuropathological level with amyloid plaques and tauopathy. The amyloid pathology is due to extracellular aggregation of soluble amyloid beta (Aβ) monomers into oligomers, fibrils, protofibrils and finally amyloid plaques. The tauopathy is mainly observed in neurons due to hyperphosphorylated tau protein aggregation. Classical hallmarks of the disease also include a neuroinflammatory response involving astrocytes and microglial cells. When they become reactive, both cell types demonstrate morphological, transcriptional and functional modifications directly impacting the pathology^1,2^.

Despite intensive research for treatments, most existing drugs only slow disease progression and alleviate symptoms. Recently, three monoclonal antibodies, Aducanumab (Adu, Biogen and Eisai), Lecanemab (Biogen and Eisai), and Donanemab (Lilly), targeting different forms of Aβ aggregates^3,4^ have been approved by the Food and Drug Administration (FDA) and, more recently, by the European Medicines Agency as new treatments for AD, the first in nearly two decades^5^. Clinical trials have demonstrated that these treatments can reduce Aβ levels and improve cognition in patients with prodromal or mild AD, with effects varying based on dosage and duration of treatment^17,20,21^. Unfortunately, those immunotherapies require a large dose infusion of antibody due to blood-brain barrier (BBB) blockage which could be associated with important side effects such as amyloid-related imaging abnormalities (ARIA), whose occurrence may dependent on the dosage^6,7^ and appears to not be limited to anti-amyloid therapies^8^. Consequently, many studies are investigating strategies to enhance the entry of therapeutic molecules into the brain, to reduce the injected peripheral dose without compromising their effectiveness.

Low-intensity focused ultrasound (LiFUS) has been extensively studied in the last decade as a non-invasive tool to modulate BBB permeability and improve drug delivery. This technology, associated with intravenous injection of microbubbles (MB), allows to create a cavitation phenomenon in contact of ultrasound that disrupt the tight junctions between endothelial cells, leading to a transient BBB opening (BBBO) demonstrated in rodent models and even in humans^9–15^. It presents several advantages for the treatment of brain disorders, mainly its non-invasiveness, allowing for repeated treatments in the case of chronic diseases. Moreover, its spatial control allows to choose the right brain location of treatment.

Importantly, LiFUS can be combined with different therapeutic agents without requiring any structural modifications to improve their brain delivery^16^. The efficacy and the safety of LiFUS-BBBO has been extensively studied and is now well demonstrated in preclinical models, including mice, rabbits, sheep, nonhuman primates, and more recently in human clinical trials^38–43^.

Interestingly, in the context of AD, LiFUS+MB demonstrated interest by itself as some studies in mouse models of the disease showed a decrease of amyloid plaques^17–21^, and even the tauopathy^22–24^. LiFUS+MB could also restore the long-term potentiation (LTP) levels in the hippocampus, essential in memory processes^25^. Additionally, most studies tend to also report an upregulation of inflammatory processes after LiFUS+MB but the time course of this response is not consistent, and underlying mechanisms remain poorly understood. Indeed, methodological heterogeneity exists across preclinical studies investigating the impact of LiFUS+MB on neuroinflammation, including differences in treatment timing, assessment windows and methods to evaluate neuroinflammatory response, as well as discrepancies in LiFUS parameters (e.g., peak negative pressure, frequency, microbubble dose, treated volume, sequence duration)^26–31^. This variability complicates direct comparisons between studies and makes it difficult to reach a clear consensus regarding the effect of LiFUS+MB on neuroinflammation. Moreover, almost all studies focused on short-term delays. Consequently, even if the safety of LiFUS+MB procedure is already investigated in human AD patients^14,32^, it appears crucial to further understand their impact on brain functions and neuroinflammatory response, particularly after repeated exposures, necessary for chronic diseases’ treatment.

In this study, we therefore used the TgF344-AD rat model of AD^33^, developing amyloid plaques, neuroinflammation and at later stage an endogenous tauopathy, to conduct an in depth analysis of the effect of a single or multiple exposure to LiFUS+MB in a large volume of the brain at different timepoints, on microglial response, Tau and amyloid load.

## Materials and methods

### Animals

Male and female wild-type (WT) and TgF344-AD (TgAD) rats carrying the human APPswedish and PS1dE9 transgenes on a Fisher 344 background^33^ were housed with food and water *ad-libitum* in a 12-hours light-dark cycle. To evaluate the acute impact of our LiFUS+MB procedure, WT or TgAD rats received one LiFUS+MB treatment at 18/19-month-old and euthanized 24h (n=6/sex) or 7 days (n=5 males, n=1 female) later. Age- and sex- matched TgAD rats receiving no ultrasound (Sham; n=6/sex) were used as control (Figure 1). To evaluate the impact of repeated LiFUS+MB sessions, TgAD rats received 4 LiFUS+MB exposure once a week at 18/19-month-old (n=4/sex) and euthanized 7 days or 6 weeks after the last treatment at 20-month-old. Control TgAD rats underwent anesthesia for 20 min each week (n=4/sex) (Figure 1). To investigate if LiFUS+MB treatment impact was dependent of the disease stage, another cohort was treated at an early stage (10-month-old) with 4 LiFUS+MB once a week (n=6/sex) and euthanized 6 weeks later. Control animals were anesthetized 20 min each week (sham-treated animals; n=6 females, 2 males) (Figure 1). Animals were randomly assigned to groups. All experimental procedures were approved by the Ethics Committee for Animal Experimentation of the Canton of Geneva, Switzerland.

**Figure 1:**
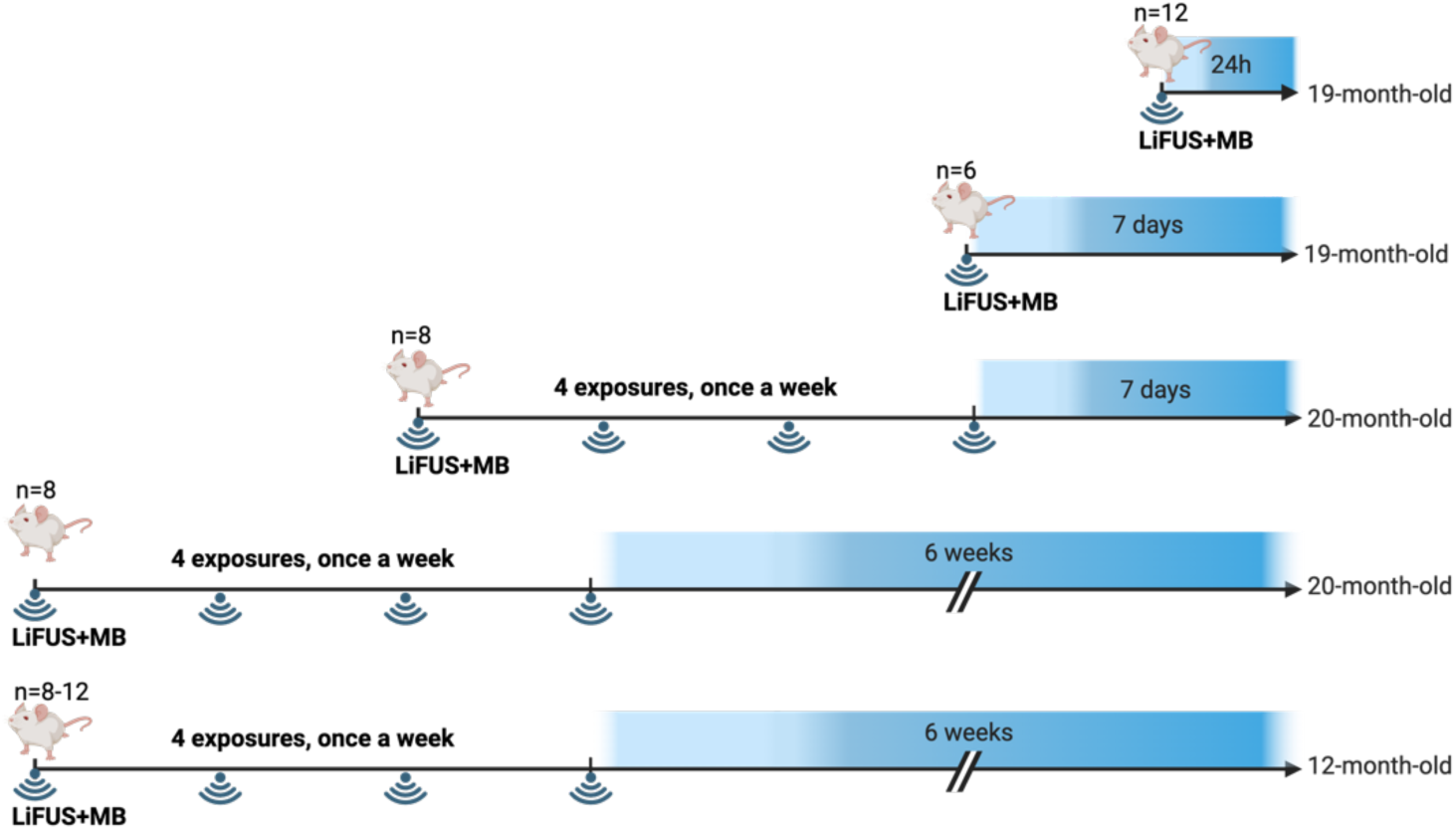
Experimental groups.

### Low-intensity Focused Ultrasound treatment

The ultrasound setup featured a 7-element annular spherical transducer (25 mm diameter, 20 mm radius of curvature, Imasonic) with a central frequency of 1.5 MHz. This transducer was driven by a 16-channels sine wave phased array generator, output power maximum 3W/channel, synchronized with a motorized XY stage (Image Guided Therapy (IGT) and comprising water degassing circuit for removing air bubbles from the water-filled balloon. The XY motorized stage allows a synchronous transmission of FUS with transducer movement (BBBop software; IGT). Acoustic pressure delivery was measured using a 200-μm calibrated hydrophone (HGL-0200, preamplifier AH-2020, Onda Corporation, USA), mounted on a micrometric 3D positioning stage and placed at the transducer’s focus in a water tank (Figure 2A). The -6 dB dimensions of the focal spot were 7 mm axially and 1 mm laterally. The acoustic transmission at 1.5 MHz through the male and female rat skulls were measured at different ages using experimental setups described by Gerstenmayer et al^34^ and taken into account to tune up the acoustic peak negative pressure (PNP) at the focal point through the skull (Figure 2B). Concretely, voltage compensation was applied according to the animal’s weight and sex to apply an estimated PNP of 450kPa *in situ* (mechanical index: 0.24).

**Figure 2:**
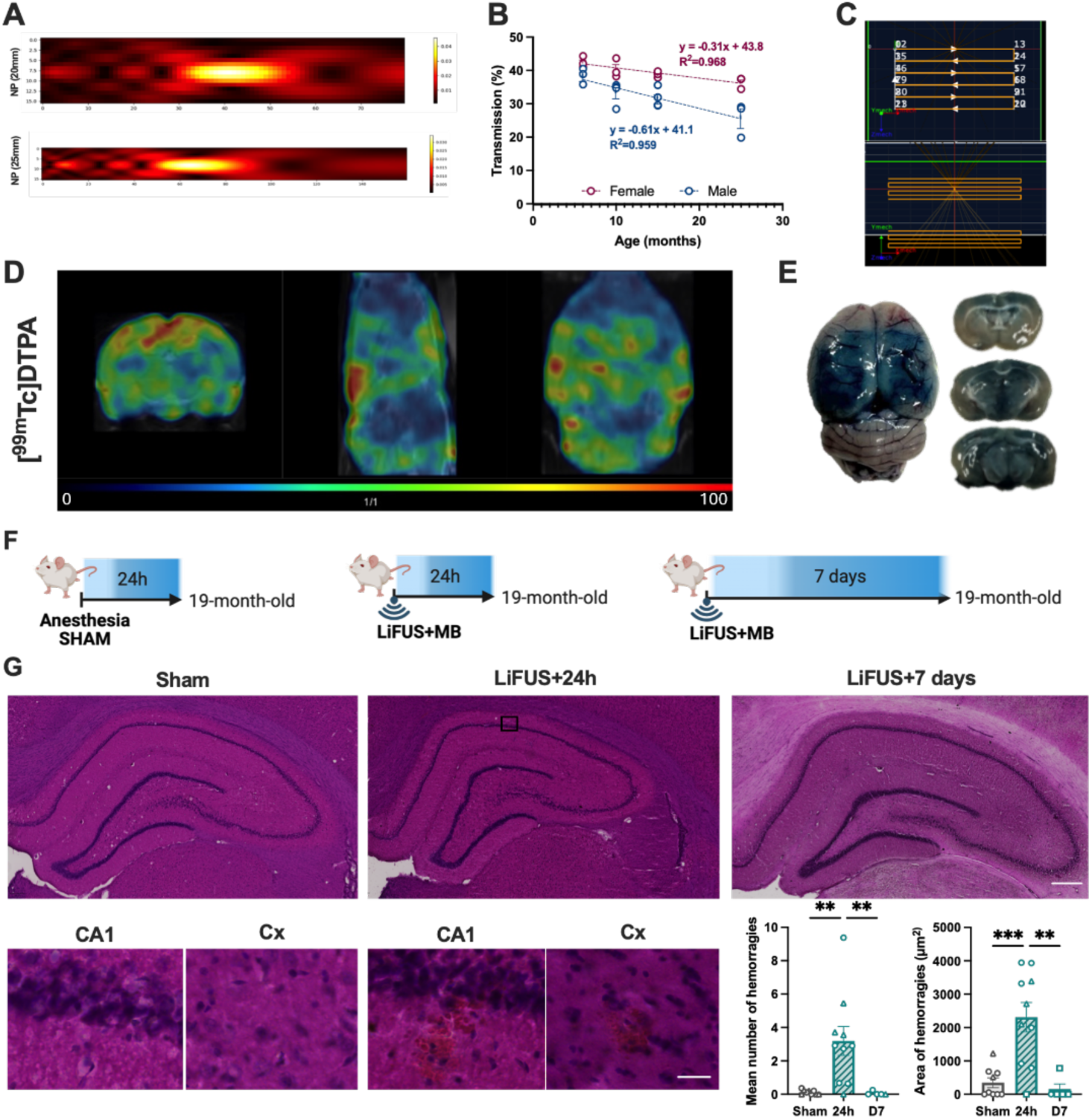
Experimental apparatus for LiFUS+MB treatment and efficient BBB opening in rats. **A)** To calibrate our apparatus for LiFUS+MB treatment, a submersible hydrophone was placed in degassed water below the transducer. The measurement of the peak negative acoustic pressure delivered by the transducer was realized at 20 (geometrical point) and 25 mm (steering). **B)** The coefficient of transmission through the skull was determined depending on the age and the sex of WT rats. **C)** A sequence with two focal points (geometrical focal point and steering point) was developed to target a large volume of the rat brain. **D, E)** [^99m^Tc]DTPA signal acquired by SPECT imaging **(D)** and Evans blue **(E)** staining in a female brain after LiFUS+MB treatment validating the BBB opening in the treated volume. **F)** Experimental design. **G)** Hematoxylin and Eosin staining 24h or 7 days after a single LIFUS+MB exposure, showing transient microhemorrhages. Scale bars = 400 µm (top), 20 µm (bottom). One-way ANOVA and Kruskal-Wallis post hoc tests. *p≤0.05, **p≤0.01. Males are represented by round points and females by triangles.

Under isoflurane anesthesia, the animals were shaved and positioned in a dedicated bed within a stereotactic frame. The transducer was coupled to the head using a latex balloon filled with deionized and degassed water, and acoustic gel was applied to the skin to ensure effective coupling with the balloon. A bolus of commercially available MB (400 µl/kg; SonoVue®, Bracco) was injected into the tail vein. Ultrasound sonication started immediately after the MB injection, using continuous waves and a mechanical raster scan. One minute later, a second bolus of MB was administered (400 µl/kg, SonoVue). For the LiFUS impact study, the transducer was moved back and forth over the posterior area of the rat brain (targeting the 2 hippocampus and cortex region above) along 6 segments at a speed of 10mm/s, covering a total surface of 12mm x 6mm (Figure 2C). Pseudo-continuous ultrasound waves were transmitted at 2 different depths (*i.e.*, geometrical focusing at 20 mm, corresponding to the geometrical point, and steering position at 25 mm from the transducer; Figure 2C) along the described trajectory for a total duration of 3.2 minutes^35^.

### SPECT imaging

For [^99m^Tc]DTPA imaging, rats (n=2) were anesthetized (3% isoflurane) and were placed in the U-SPECT-II scanner (MiLabs). A tail-vein injection of the radiotracer [^99m^Tc]DTPA (50 MBq) was performed and a 30min SPECT scan was acquired 30 min after injection. Animals were immediately euthanized after SPECT imaging.

#### Evans blue

An Evans blue 4% (2 ml/Kg) solution was injected in the tail vein of animals (n=2), 5 min after the LiFUS+MB sequence. The animal was euthanized by an injection of pentobarbital 60 min later. The brain was then dissected in 1mm slices in a brain matrix to assess the surface and depth of the BBB opening.

#### Histology

Twenty-four hours after the LiFUS+MB treatment, TgAD rats were euthanized under anesthesia (2% isoflurane) by intracardiac perfusion with a saline solution. One hemisphere was postfixed in a 4% paraformaldehyde solution for 24h and cryoprotected with a sucrose gradient (10 to 20%) before being frozen in isopentane. Serial brain sections (35 µm) were realized with a cryostat and stored in an anti-freeze solution at -20°C until used for histology. The other hemisphere was rapidly dissected to isolate the hippocampus and surrounding cortex, manually homogenized together, and snap frozen in liquid nitrogen for RT-qPCR measures or biochemistry. After washes in PBS0.1M, slices were mounted on slides and let dry O/N. Slices were rehydrated in PBS0.1M and were incubated in a hematoxylin and Eosin (H&E) solution to evidence potential tissue damages. For inflammation markers’ analysis, slices were incubated O/N at 4°C with the following primary antibodies: anti-GFAP-Cy3 (1/1000, Mouse; Sigma) or anti-IBA1 (1/500, Rabbit; Wako) in 1% BSA/PBS0.1M/0.3% Triton X-100. After 3 washes in PBS0.1M, slices were incubated for 1h at RT in the appropriate secondary antibody (1/250, Alexa Fluor) in 1% BSA/PBS0.1M/0.3% Triton X-100. Finally, slices were rinsed before being coverslipped with Fluorosave^TM^ (Calbiochem).

#### Histological image analysis

Images were acquired at 10x or 20x magnification using the Axioscan.Z1 scanner (Zeiss) and analyzed with the QuPath software^36^. For H&E, microhemorrhages were manually delimited and the mean number of events per slices and the mean area (µm^2^) of each microhemorrhage was calculated. For immunostaining, the hippocampus and the surrounding cortex were manually delineated. An intensity threshold, consistent across all subregions, was applied to quantify the percentage of the region of interest (ROI) that was positively stained with GFAP, or IBA1.

#### Protein extraction

Animals were euthanized under anesthesia (2% isoflurane) by intracardiac perfusion with a saline solution. Both hemispheres were dissected to collect the hippocampus and the adjacent cortex for biochemistry or flow cytometry (see details below). Proteins were extracted in a Triton X100 lysis buffer [50 mM Tris-HCl pH = 7.4, 150 mM NaCl, 1% Triton X-100 with 1x protease and phosphatase inhibitors (Pierce); 500 μl], centrifuged at 20,000 g for 20 min at 4 °C. The supernatant contains Triton X100 (Tx)-soluble proteins. The pellet was re-suspended in a Guanidine lysis buffer [50 mM Tris-HCl pH = 8, 5 M Guanidine HCl with 1x protease and phosphatase inhibitors (Pierce); 250 μl], incubated for 3h on ice and centrifuged at 20,000 g for 20 min at 4 °C. The supernatant contained Guanidine (Gua)-soluble proteins. The total protein concentration was determined using the BCA test (Pierce).

#### ELISA tests

Aβ peptides were measured using the human Aβ40 ELISA test (Life Technologies), ultrasensitive human Aβ42 ELISA test (Life Technologies) and human Aβ42 ELISA test (Life Technologies). PT231 levels were measured using the human Tau (phospho) pT231 ELISA kit (Life Technologies). If necessary, Tx-soluble samples were pre-diluted in the kit diluent, and Gua-soluble samples were first diluted in Gua-lysis buffer and then diluted at 1/500 in the kit diluent. Manufacturer’s protocols were followed. Absorbances were measured at 450 nm using the SpectraMax iD3 plate reader (Molecular devices). The absence of amyloid in WT was validated for all forms. Aβ concentrations were determined using a 4S parameters analysis and were normalized to the total protein quantity (µg proteins).

#### [^125^I]CLINDE synthesis

[125I]CLINDE were labeled as previously described^37^. Briefly, the CLINDE tributyltin precursor (100 µg) was resuspended in 10µl of Hexafluoro-2-propanol (HFIP). The precursor was then incubated with 3mg of Sodium acetate, 95µl of N-Chlorosuccinimide (at 0.016M in HFIP) and 10µl of [^125^I]NaI (10MBq, PerkinElmer) for 25 min at RT. Reactions were then diluted in 50% acetonitrile (ACN) in water (350µl), injected onto a reversed-phase column (Bondclone C18) and [^125^I]CLINDE, a TSPO specific radiotracer, was isolated by a linear gradient HPLC run (5% to 95% ACN in 7mM H3PO4, 10 min). At the end, ACN was evaporated and [^125^I]CLINDE was dissolved in saline.

#### FACS-RTT

The FACS-RTT technique was previously developed by authors to evaluate the radioligand binding in specific cell types^38^. Hippocampal and cortical tissues were collected in cold HBSS before a mechanical and enzymatic dissociation followed by a myelin depletion step (Miltenyi biotec). Cells were then suspended in 0.1 M PBS/ 1 mM EDTA/ 1% of BSA buffer and incubated with [^125^I]CLINDE (1.4MBq/sample) for 15 min on ice. After a washing step, cells were treated with an anti-rat CD32 (1/100; BD Bioscience) for 5 min on ice to block Fc-mediated adherence of antibodies. After a centrifugation at 350 g, 5 min at 4°C, cells were resuspended with a solution of the primary antibody, PerCP/Cyanine5.5 anti-rat CD11b/c (1/100; 201820, Biolegend), diluted in 0.1 M PBS/ 1 mM EDTA/ 1% of BSA buffer for 20 min on ice. After washing and centrifugation at 350 g, 5 min at 4°C steps, cells were suspended in 400 µl of 0.1 M PBS for cell sorting. Unstained and single stain cells were used to identify positive and negative cells for each antibody, to identify auto-fluorescent cells, to determine the values of compensation (to correct for interference between fluorochromes) and to draw sorting gates for each antibody. Cells were first sorted based on their forward and side scatter from all detected events. Dead cells were excluded using DAPI staining before cell sorting. Microglia cells were sorted based on their fluorescence for PerCP/Cy5.5-CD11b. Cell numbers were determined during cell sorting on a BD FACSAria™ Fusion Flow Cytometer (BD Biosciences). Radioactive concentrations in sorted microglia were measured on an automatic *γ* counting system (Wizard 3”, PerkinElmer). Data are expressed in count per minute (CPM)/g of tissue or CPM/number of cells for each animal.

#### RNA extraction and qPCR

Samples (sorted cells or tissue homogenates) were placed into 400 µl of Trizol before storage at -80°C. The day of the extraction, samples were left at RT for 5 min. Then, 100 µl of chloroform was added for 3 min. After a vortex and centrifugation at 12,000 g for 15 min at 4°C, the aqueous phase was collected and 1 volume of ethanol 70% was added. The whole volume was then transferred onto RNeasy columns (RNeasy micro kit, Qiagen) and the manufacturer’s protocol was followed. RNA was eluted with 15 µl of nuclease free water. Superscript IV VILO synthesis kit (Invitrogen) was used to synthesized cDNA as described by the manufacturer. Samples were diluted at 1/10 in H2O and mixed with 250 nM of primers and PowerUp SYBR Green Master Mix (Applied Biosystems) for qPCR. The following sequences of primers were used: *Aif1-F:* CAGAGCAAGGATTTGCAGGGA*, Aif1-R:* CAAACTCCATGTACTTCGTCTTG*; ApoE-F: CTGAACCGCTTCTGGGATTACCTG; ApoE-R: CATAGTGTCCTCCATCAGTGCCGTC; C1qa-F : TCAGCTATTCGGCAGAACCC, C1qa-R : GGGTCACCAAAGCCAGAAAATG; C3-F: GACCTGCGACTGCCCTACTCT, C3-R: CTGATGAAGTGGTTGAAGACG; Clusterin-F:* AGCGCACTGGAGCCAAG, *Clusterin-R:* TCAGAGACCTCCTGCTCTCC; *Cx3cr1-F:* TCTTCACGTTCGGTCTGGTG*, Cx3cr1-R:* GGCAAAGTGGCCACAAAGAG*; Gfap-F: TTGACCTGCGACCTTGAGTC, Gfap-R: GAGTGCCTCCTGGTAACTCG; Glut1-F:* GCATCTTCGAGAAGGCAGGT*, Glut1-R:* AACAGCGACACCACAGTGAA*; Il1β-F:* CACACTAGCAGGTCGTCATCATC, *Il1β-R:* ATGAGAGCATCCAGCTTCAAATC; *Il4-F:* AGACGTCCTTACGGCAACAA, *Il4-R:* CACCGAGAACCCCAGACTTG; *Il6-F:* AGAGACTTCCAGCCAGTTGC, *Il6-R:* AGTCTCCTCTCCGGACTTGT; *Mbp-F:*TCTCAGACCGCCTCAGAAGA*, Mcu-F: ACAGTTTGGCATTTTGGCCC, Mcu-R: ATTCCTGGCGCGTCATTACA; Mbp-R:* TGTGCTTGGAGTCTGTCACC*; Mct1-F:*GCTGCTTCTGTTGTTGCGAA*, Mct1-R:* AAATCCAAAGACTCCCGCGT*; Mct4-F:* CGATACTTCAACAAGCGCCG, *Mct4-R:* CAGTGCACAAAGGAACACGG*; Mertk-F:* CAGTACCGAAGTCCATCCCC, *Mertk-R:* CCGAGCAGCTAAATCCCTGT; *NeuN-F:* GAGTCTATGCGGCTGCTGAT, *NeuN-R:* CAGAGGCTAGCCATGGTTCC*; Ppia-F:* ATGGCAAATGCTGGACCAAA, *Ppia-R:* GCCTTCTTTCACCTTCCCAAA; *P2y12r-F:* AACGCCTGCCTTGATCCATT*, P2y12r-R:* TACATTGGGGTCTCCTCGCT*; Rantes-F: GCAGTCGTCTTTGTCACTCG, Rantes-R: GGAGTAGGGGGTTGCTCAGT; Stat3-F:* CAACTTCAGACCCGCCAACA, *Stat3-R:* GCCGCTCCACCACGAAGG; *Tgfβ-F:* CCTGGAAAGGGCTCAACAC, *Tgfβ-R:* CAGTTCTTCTCTGTGGAGCTGA; *Tnfα-F:* CAGAGCAATGACTCCAAAGTA, *Tnfα-R:* CAAGAGCCCTTGCCCTAA*; Trem2-F:* CCTGTGGGTCACCTCTAACC*, Trem-R:* GGCCAGGAGGAGAAGAATGG; *Tspo-F:* GCTGCCCGCTTGCTGTATCCT*, Tspo-R:* CCCTCGCCGACCAGAGTTATCA*; Vdac1-F:* GTCACCGCCTCCGAGAACAT*, Vdac1-R:* CGTAGCCCTTGGTGAAGACA; *Vimentin-F*: CAGTCACTCACCTGCGAAGT, *Vimentin-R*: AGTTAGCAGCTTCAAGGGCA. Expression levels of genes of interest were normalized to the abundance of *Ppia* gene with the ΔCt method.

#### Statistics

Analyses were performed in blind conditions and normality of residues was assessed with the Shapiro-Wilks test. An unpaired *t-*test was performed if the normality conditions were respected, otherwise, a Mann-Whitney test was performed. p-values were then adjusted for multiple comparisons using the Benjamini-Hochberg False Discovery Rate (FDR) procedure. Genes with an adjusted p-value (q-value) less than 0.05 were considered statistically significant. For H&E analysis, one-way ANOVA and Tukey’s multiple comparisons test or Kruskal-Wallis followed by a Dunn’s multiple comparisons test were used depending on normality of residues. Outliers were identified using the ROUT method (Maximal false discovery rate = 1%). All analyses were performed on GraphPad Prism 10. Statistical details are provided only if significance was reached.

## Results

Multiple studies described a safe BBB opening in rodents with an *in situ* peak negative acoustic pressure (PNP) set at 450 kPa^31,39^. Consequently, our LiFUS apparatus was first calibrated to determine the amplitude necessary to reach a maximum of 450 kPa at the geometrical focal point (focal depth at 20 mm) or at the steering point (focal depth at 25 mm) (Figure 2A and Supplemental figure 1). The impact of skull thickness on ultrasound transmission was also measured in our TgAD rat model. As sex and age are two factors influencing skull thickness, the coefficient of transmission (%) of ultrasound through the skull was measured for males and females at different ages (6, 10, 16, and 25-month-old; Figure 2B). It is important to note that animals begin to lose weight after a certain age. Therefore, considering the age of the animals rather than their weight appears more relevant, particularly when using aged animal models for studying degenerative diseases. Once LiFUS parameters were determined to reproducibly apply an acoustic pressure of 450 kPa, a sequence to target a large volume of the rat brain (about 640 mm^3^ considering the size of the focal spot and the trajectory) was developed (Figure 2C). The efficient BBB opening was validated by SPECT imaging using [^99m^Tc]DTPA (Figure 2D) or an intravenous injection of Evans blue (Figure 2E), which only penetrate the brain in the presence of an alteration of the BBB. The safety of the procedure was then assessed using H&E staining to detect potential microhemorrhages. At T+24h, microhemorrhages were detectable in the entire targeted area (cortex, striatum, hippocampus or thalamus), that were completely resorbed 7 days post treatment (Figures 2G; Number of hemorrhages: Kruskal-Wallis *p=*0.0012, Dunn’s post hoc test: Sham vs 24h *p=*0.0084, Sham vs D7 *p>*0.9999, 24h vs D7 *p=*0.0052; Area of hemorrhages: One-way ANOVA *p=*0.0002, Tukey’s post hoc test: Sham vs 24h *p=*0.0006, Sham vs D7 *p=*0.9335, 24h vs D7 *p=*0.0015).

Then, the effect of a single sonication on amyloid plaques, Tau and inflammatory cells was investigated (Figure 3A). The concentration of soluble/poorly aggregated forms of Aβ40 (Tx-Aβ40) and highly aggregated forms (Gua-Aβ40) were not impacted 24h after sonication (Figure 3B), contrary to the highly aggregated Aβ42 that was significantly reduced (Mann-Whitney test *p*=0.0226; Figure 3C). This reduction was confirmed with Methoxy-X04 staining, a derivative conjugate of Congo Red labeling dense-core amyloid plaques, in the hippocampus (Supplemental figure 2). No significant effect was observed on pTau231 concentration (Figure 3D). The percentage of IBA1^+^ area remained unchanged in hippocampal subregions and in the surrounding cortex 24h after LiFUS exposure (Figure 3E-G).

**Figure 3:**
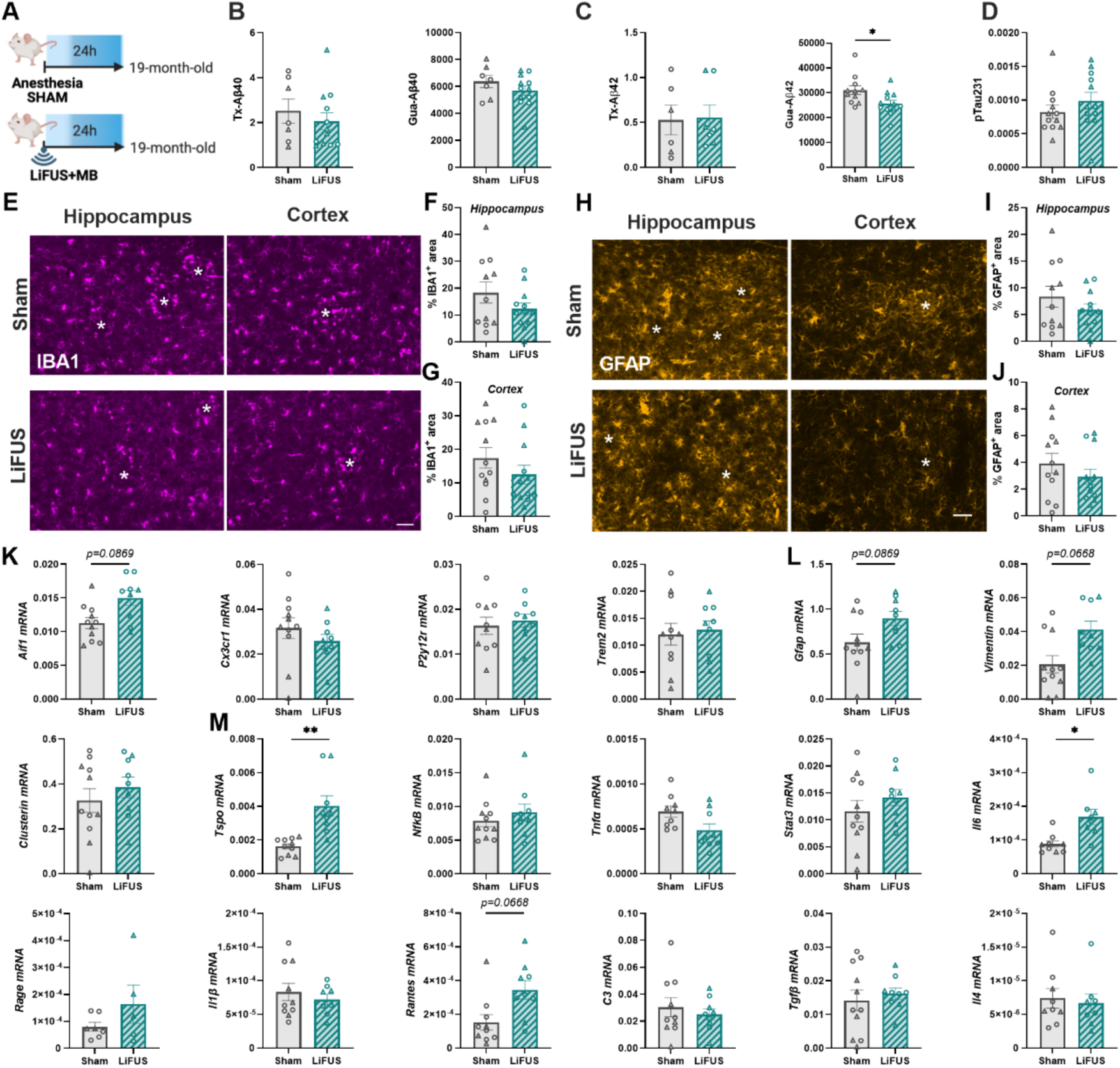
A single LiFUS+MB exposure slightly modulates astrocyte and microglial reactivity 24h post-treatment in TgAD rats. **A)** Experimental design. **B, C)** Triton (Tx)-soluble and Guanidine (Gua)-soluble forms of Aβ40 (**B**) and Aβ42 (**C**) were measured by ELISA in the hippocampus + adjacent part of the cortex in sham-treated TgAD rats and LIFUS+MB-treated. **D)** pTau231 levels were quantified by ELISA 24h after a single LIFUS+MB exposure. Unpaired student *t* test or Mann-Whitney test. **E)** Representative images of IBA1 (magenta), a microglial marker, in the hippocampus and the cortex of sham-treated TgAD rats and LiFUS+MB-treated TgAD rats. Stars represent amyloid plaque localization. Scale bar = 50 µm. **F, G)** Quantification of the % IBA1^+^ area in both regions 24h after a single LIFUS+MB exposure. **H)** Representative images of GFAP (orange), an astrocytic marker, in the hippocampus and the cortex of sham-treated TgAD rats and LiFUS+MB-treated TgAD rats. Stars represent amyloid plaque localization. Scale bar = 50 µm. **I, J)** Quantification of the % GFAP^+^ area in both regions 24h after LIFUS+MB exposure. Unpaired student *t* test. **K-M)** mRNA levels of microglial (**K**), astrocytic (**L**) and inflammatory (**M**) genes in the hippocampus and surrounding cortex homogenate in sham vs LiFUS+MB-treated groups 24h after LiFUS exposure. FDR multiple correction test. *p-adj≤0.05, **p-adj≤0.01. Males are represented by round points and females by triangles.

Similarly, the percentage of GFAP^+^ cells was not modified in the treated group (Figure 3H-J), suggesting no modulation of neuroinflammation at the protein levels at an advanced disease stage. To go further, the mRNA expression of several glial and inflammatory markers in the brain (hippocampus and surrounding cortex) of sham- and LiFUS+MB-treated rats was measured. Among microglial gene expression quantified, only *Aif1*, the gene encoding IBA1, tended to be overexpressed 24h after LiFUS (FDR *p-adj=*0.0869; Figure 3K). Moreover, among astrocytic markers, *Gfap* (FDR *p-adj=*0.0869) and *Vimentin* (FDR *p-adj=*0.0668) mRNA levels also tended to be increased (Figure 3L). Other microglial and astrocytic gene expression remained unchanged (Figure 3K, L). To deepen characterize the inflammatory response at the transcriptional level, the expression of pro-and anti-inflammatory gene levels was quantified (Figure 3M). The 18kDa translocator protein (TSPO) is a classical marker of neuroinflammation known to be overexpressed in glial cells of animal models and AD patients’ brain, even before the appearance of classical neurochemical markers and cognitive symptoms of AD^40–43^. Interestingly, an increase in *Tspo* mRNA levels was measured after LiFUS+MB (FDR *p-adj=*0.0025). Furthermore, the interleukin 6 *(Il6;* FDR *p-adj =*0.0273) and chemokine *Rantes/Ccl5* (FDR *p-adj=*0.0668) gene expression were upregulated in LiFUS+MB-treated group compared to sham-treated animals. These changes suggest a slight inflammatory response to a single LiFUS+MB exposure 24h post-treatment at the transcriptomic level. No difference between males and females were observed.

In the context of AD, a chronic treatment protocol is more translationally relevant (Figure 4A). Even if the safety of our LiFUS procedure was validated with a complete resorption of microhemorrhages 7 days after a single treatment (interval chosen between two LiFUS+MB sessions), we first confirmed that multiple exposures did not increase this recovery time using H&E staining (*data not shown)*. Then, the effect of multiple sonication (*i.e.*, a total of 4 sessions repeated once a week) on AD hallmarks 7 days after repeated LiFUS treatments was assessed. As after a single sonication, Aβ40 levels were not impacted (Figure 4B). Surprisingly, a significant increase of highly aggregated forms of Aβ42 was measured (Mann-Whitney test *p=*0.0059) in LiFUS-treated animals, without effect on soluble or poorly aggregated peptides (Figure 4C). The concentration of pT231 was also increased after chronic LiFUS sonication (Unpaired student *t* test *p=*0.0009; Figure 4D). The expression of several microglial genes was altered (Figure 4E), with increased expression of *Aif1* (FDR *p-adj*=0.0269) and *P2yr12r* (FDR *p-adj*=0.0179), and decreased expression of *Cx3cr1* (FDR *p-adj*=0.0111). *Trem2* mRNA levels also tended to be decreased (FDR *p-adj*= 0.0548). These changes suggest an impact of chronic LiFUS treatment on microglial inflammatory status. By contrast, astrocytic gene expression remained unchanged (Figure 4F), with the exception of *Clusterin* mRNA, which was increased (FDR *p-adj*=0.0111). As observed after a single LiFUS exposure, *Tspo* mRNA levels were also increased 7 days after repeated treatment (FDR *p-adj*=0.0209, Figure 4G). Among the inflammatory genes, an increase of *Rantes/Ccl5* (FDR *p-adj=*0.0468) and a decrease of the *C3 complement* (FDR *p-adj=*0.0111) mRNA levels were measured in LiFUS+MB-treated group. *Stat3* gene expression also tended to be reduced in this group (FDR *p-adj*=0.0896). Overall, these results suggested that a short-term inflammatory response persists 7 days after chronic LiFUS+MB treatment, with a predominant impact on microglial cells.

**Figure 4:**
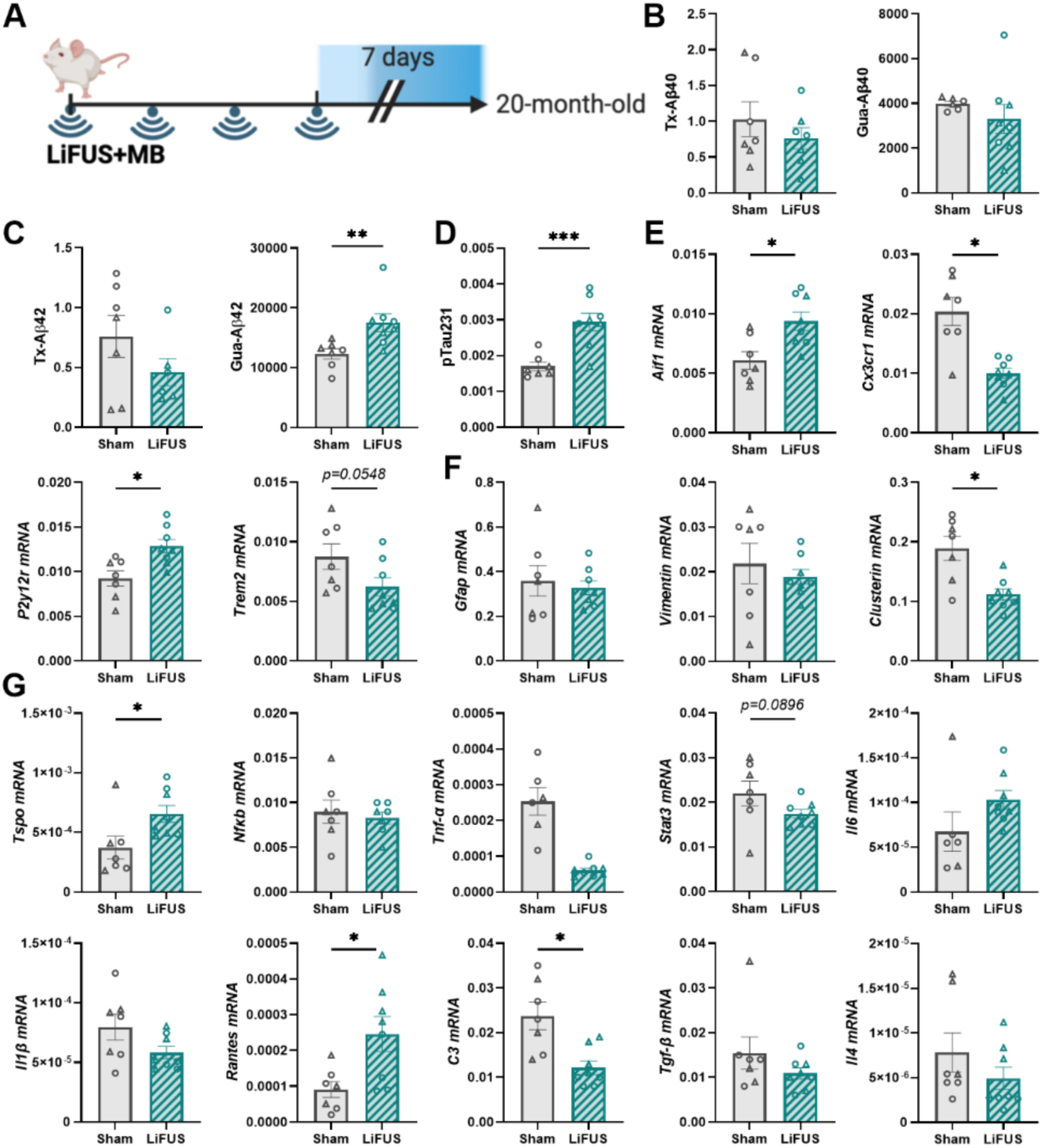
Repeated LiFUS+MB exposures induce a short-term inflammatory response 7 days post-treatment in old TgAD rats. **A)** Experimental design. **B-C)** Triton (Tx)-soluble and Guanidine (Gua)-soluble forms of Aβ40 (**B**) and Aβ42 (**C**) were measured by ELISA in the hippocampus + adjacent part of the cortex in sham-treated TgAD rats and LIFUS+MB-treated TgAD rats. **D)** pTau231 levels were also quantified by ELISA in old TgAD rats. Unpaired student *t* test or Mann-Whitney test. **E-G)** mRNA levels of microglial (**E**), astrocytic (**F**) and inflammatory (**G**) genes in the hippocampus and surrounding cortex homogenate in sham vs LiFUS+MB-treated groups 7 days after the last LiFUS exposure. FDR multiple correction test. *p-adj≤0.05, ***p≤0.001. Males are represented by round points and females by triangles.

Microglia are highly adaptive cells that functionally respond to changes in their environment, and seems to be particularly impacted by LiFUS, as already described^44^. Consequently, in another cohort, the effect of repeated LiFUS+MB treatment on neuroinflammation was investigated at long term (6 weeks post-treatment, Figure 5A), with a focus on microglial cells. AD hallmarks were first quantified as for previous cohorts. Repeated LiFUS exposures reduced both forms of Aβ40 (Tx-Aβ40: Unpaired student *t* test *p=0.0648,* Gua-Aβ40: Unpaired student *t* test *p=0.0138;* Figure 5B), without influence Aβ42 (Figure 5C) or pT231 levels (Figure 5D). The effect of repeated LiFUS treatment was then measured at the cellular level on sorted microglia (CD11b^+^) from hippocampus and cortex region of sham-treated and LiFUS+MB-treated TgAD rat brain. Sorted cells highly expressed genes known as microglia markers (*Aif1* and *Cx3cr1*) whereas the other cell type markers were weakly expressed (*Gfap*, astrocytes; *Neun*, neurons; *Mbp*, oligodendrocytes), validating the efficient sorting of microglia (Figure 5E). The inflammatory state of microglia was first investigated (Figure 5F). Among the genes evaluated, only *Tspo* mRNA expression tended to be reduced at the long-term following repeated LiFUS+MB treatment (FDR *p-adj=*0.0519), in contrast to what was observed in the short-term. To better understand the changes induced by a repeated LiFUS+MB treatment on microglia profile, other important microglial functions were also explored. As TSPO is expressed in the outer mitochondrial membrane, the expression of genes involved in mitochondrial functions was measured (Figure 5G). However, the mRNA level of the voltage dependent anion channel 1 (*Vdac1*) and the mitochondrial calcium uniporter (*Mcu*) genes were not affected by repeated LiFUS+MB exposures. Another important cell function known to be impacted in AD is the metabolism, including in microglia cells^45^. No difference between groups was observed for glucose transporter 1 (*Glut1*), monocarboxylate transporter-1 (*Mct1;* playing a role in the lactate and ketone bodies transport), *Mct4* and apolipoprotein E (*ApoE*; involved in cholesterol metabolism) gene expression (Figure 5H). Similarly, repeated LiFUS+MB exposure did not alter the expression of genes associated with microglial phagocytosis (Figure 5I). In AD, microglial cells are known to participate in phagocytic processes^45^, notably through complement C1q A chain (C1qa)–dependent synaptic pruning^46^, as well as through the clearance of Aβ via specific receptors, including triggering receptor expressed on myeloid cells 2 (TREM2)^47,48^ and myeloid epithelial reproductive tyrosine kinase (MERTK)^49^. However, none of these phagocytosis-related pathways appeared to be involved in the reduction of Aβ40 levels in our conditions. Overall, these results suggest that repeated LiFUS+MB exposures did not markedly influence the long-term microglial cell profile and function when applied at an advanced stage of the disease.

**Figure 5:**
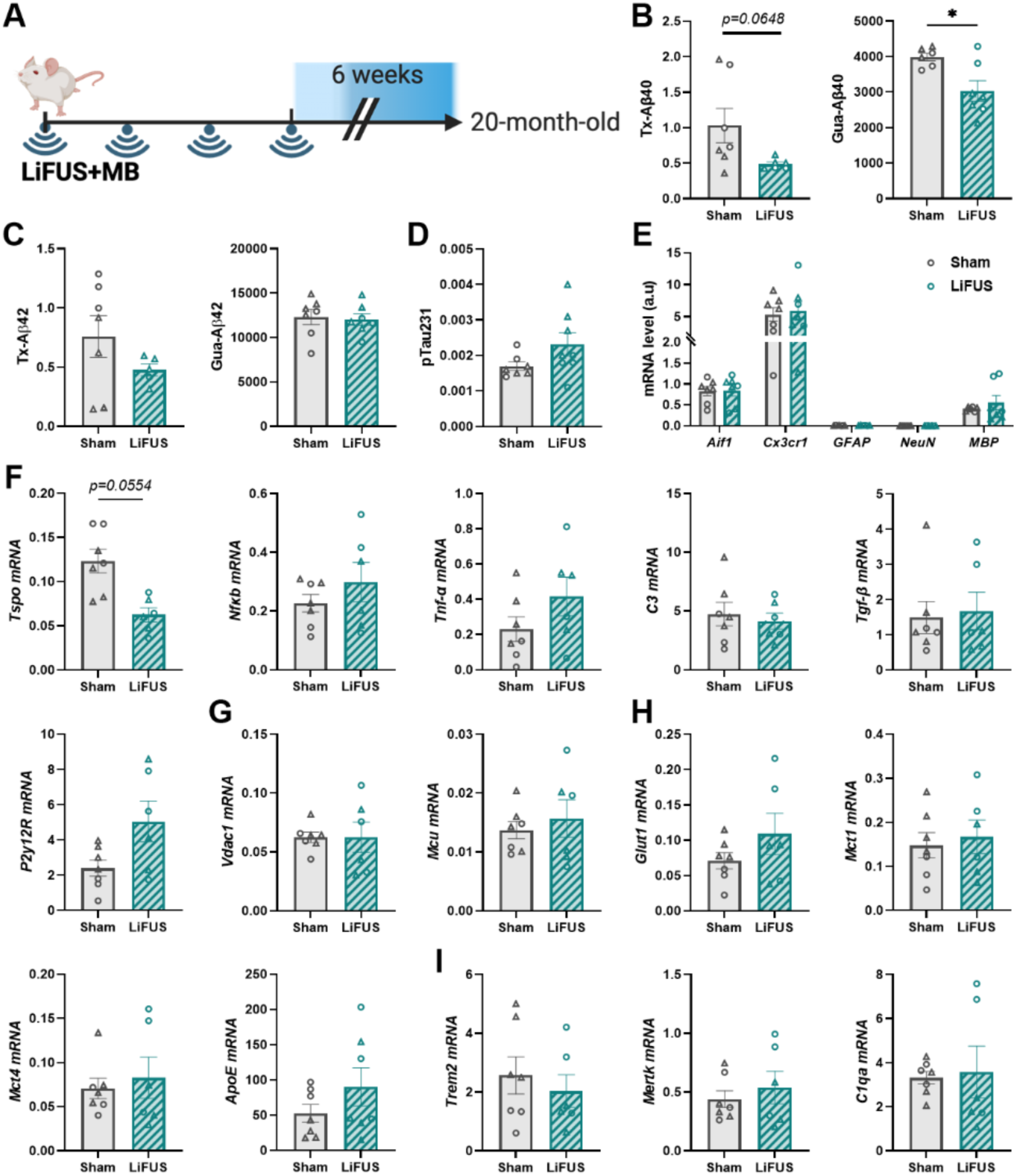
Repeated LiFUS+MB exposures does not impact microglial functions 6 weeks post-treatment in old TgAD rats. **A)** Experimental design. **B-C)** Triton (Tx)-soluble and Guanidine (Gua)-soluble forms of Aβ40 (**B**) and Aβ42 (**C**) were measured by ELISA in the hippocampus + adjacent part of the cortex in sham-treated TgAD rats and LIFUS+MB-treated TgAD rats. **D)** pTau231 levels were also quantified by ELISA in old TgAD rats. Unpaired student *t* test. *p-adj≤0.05. **E)** Sorted cells expressed microglia genes (*Aif1* and *Cx3cr1*) but did not express astrocyte (*Gfap*), neuron (*NeuN*) and oligodendrocyte (*Mbp*) specific genes. **F-I)** The mRNA expression of different markers of inflammation (**F**), mitochondria (**G**), metabolism (**H**) and phagocytosis (**I**) in microglia was not modified by repeated LiFUS+MB treatment at long-term. FDR multiple correction test. Males are represented by round points and females by triangles.

Finally, to evaluate if the stage of the disease influences the effect of LIFUS+MB on AD hallmarks, those measurements were repeated in pre-symptomatic TgAD rats of 12-months-old (Figure 6A). Repeated LiFUS treatment did not impact amyloid load or pT231 levels in younger animals (Figure 6B-D). After validation of the efficient cell sorting of microglial cells (Figure 6E), the impact of this treatment on microglia cells was evaluated. The LiFUS+MB treatment did not impact the expression of cell type markers and the proportion of sorted microglia (Figure 6E, F). As *Tspo* was one of the only genes modulated by LiFUS on aged rats, the inflammatory state of microglia with this marker was then investigated by measuring in microglia sorted cells the *in vitro* [^125^I]CLINDE binding, a radioligand of TSPO. Interestingly, a decrease in TSPO levels was observed in the total parenchyma before cell sorting (Mann-Whitney test *p=0.0314;* Figure 6G), as well as in sorted microglia (*t-*test *p<0.0001;* Figure 6H) in LiFUS+MB-treated rats, suggesting that TSPO downregulation in microglia could explain in part the total downregulation of TSPO observed after repeated LiFUS treatment. This decrease in microglia was then validated at the mRNA level. *Tspo* gene expression also tended to be reduced (FDR *p-adj=*0.0694). This reduction was accompanied by an increase expression of *Tnfα (*FDR *p-adj=*0.0279), a pro-inflammatory marker, and a decrease of the *P2y12R g*ene expression (FDR *p-adj=*0.0312) after LiFUS+MB treatment (Figure 6I). P2Y12R is described as one of the most specific microglia markers and a loss of its expression in microglia has been reported to be associated with pro-inflammatory microglia in several diseases linked to neuroinflammation, such as AD^50,51^. Moreover, the transforming growth factor beta (*Tgfβ*) mRNA levels tended to be increased (FDR *p-adj=*0.0506, Figure 6I). No changes in the mRNA expression of others inflammatory genes, including the nuclear factor-kappa B (NFκB), the receptor TREM2 and the C3 complement protein, were detected (Figure 6I). Overall, a repeated treatment of LiFUS+MB applied at an early stage of the disease slightly impacted the inflammatory status of microglia long term after a chronic treatment.

**Figure 6:**
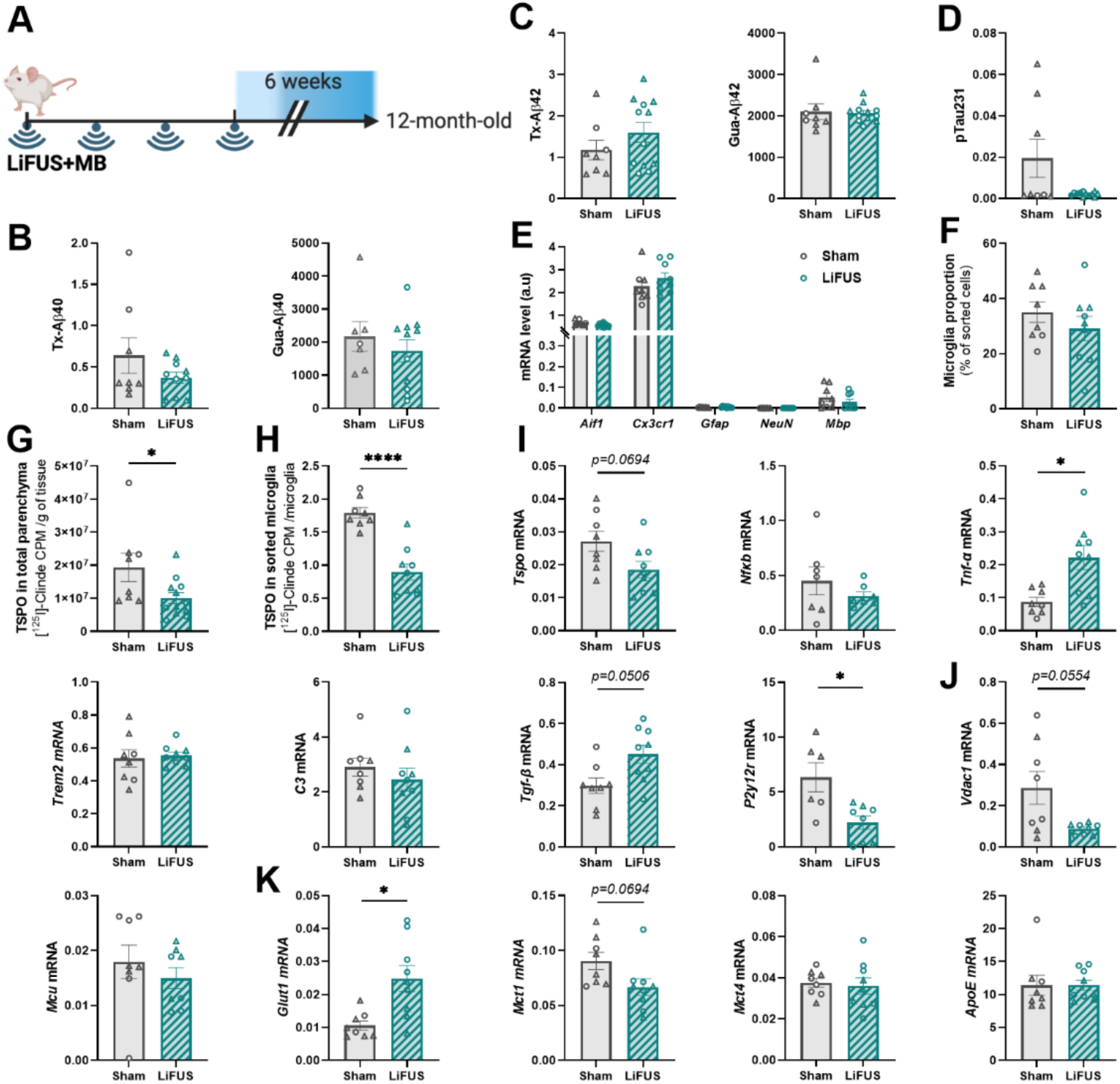
Repeated LiFUS+MB exposures impact microglial functions 6 weeks post-treatment in young TgAD rats. **A)** Experimental design. **B-C)** Triton (Tx)-soluble and Guanidine (Gua)-soluble forms of Aβ40 (**B**) and Aβ42 (**C**) were measured by ELISA in the hippocampus + adjacent part of the cortex in sham-treated TgAD rats and LIFUS+MB-treated TgAD rats. **D)** pTau231 levels were also quantified by ELISA in old TgAD rats. Unpaired student *t* test or Mann-Whitney test. **E)** Sorted cells expressed microglia genes (*Aif1* and *Cx3cr1*) but did not express astrocyte (*Gfap*), neuron (*NeuN*) and oligodendrocyte (*Mbp*) specific genes. **F)** The percentage of sorted microglia was not modified by LiFUS+MB treatment. **G, H)** *In vitro* quantification of [^125^I]CLINDE binding (count per minute, CPM), a TSPO radioligand. TSPO levels decreased in total parenchyma (hippocampus+ cortex, **G**) and in sorted microglial cells (**H**) after repeated treatment of LiFUS+MB. Unpaired student *t* test. **I-K)** The mRNA expression of different markers of inflammation (**I**), mitochondria (**J**) and metabolism (**K**) in microglia was modified by repeated LiFUS+MB treatment. FDR multiple correction test. *p≤0.05, **p≤0.01, ***p≤0.001, ****p≤0.0001. Males are represented by round points and females by triangles.

Regarding the mitochondrial function, the *Vdac1* gene expression tended to be decreased in the microglia of LiFUS+MB-treated TgAD rats compared to the sham-treated group (FDR *p-adj=*0.0554, Figure 6J), without any modulation of *Mcu* gene levels. Interestingly, regarding the metabolic functions, an increase of *Glut1* expression in sorted microglia was observed after LiFUS (FDR *p-adj=*0.0279, Figure 6K). On the contrary, the mRNA level of *Mct1* tended to be decreased in microglia of the LiFUS+MB-treated group (FDR *p-adj=*0.0694, Figure 6K). As observed at advanced pathology stage, no difference in the mRNA expression of *Mct4* and *ApoE* has been measured after the LiFUS+MB treatment (Figure 6K). These results suggest a moderate long-term adaptation of microglia following repeated LiFUS+MB treatment at early stages of AD pathology by modifications in their inflammatory status and associated with some changes in the expression of genes involved in key microglia functions in the context of AD.

## Discussion

For the development of our LiFUS+MB experimental procedure, the measure of skull thickness demonstrated a clear difference between sex and age. Both parameters are essential to consider when setting up the experimental conditions for LiFUS exposure in rats. Moreover, the age appears more relevant than weight, particularly when working with old animals, as animals lose weight after a certain age. With the adjustment of our LiFUS+MB protocol based on sex and age, our results showed that the treatment did not affect male and female rats differently. Some brain hemorrhages were observed 24h after treatment, independently of the depth of the brain region observed, but were completely resorbed 7 days later. Importantly, a repeated treatment did not amplify this side effect as no hemorrhage was observed 7 days after 4 weekly exposures.

To first investigate the impact of a single exposure to LiFUS+MB on neuroinflammation, a large volume of the brain of TgAD rats was treated and analyzed 24h post-treatment. Interestingly, a single session of LiFUS+MB led to an overexpression of inflammatory genes including *Aif1* (microglia specific), *Gfap* and *Vimentin* (astrocyte specific), *Tspo, Il6* and *Rantes* (pro-inflammatory cytokines) 24h after treatment. However, any change was observed at the protein levels when the % area positively stained for IBA1 or GFAP was quantified. The majority of studies tends to agree on a pro-inflammatory effect of LiFUS+MB but the time course of microglial and astrocyte reactivity diverge between studies^17,26–30^. We cannot exclude that, in our study, the neuroinflammatory response is too slight to be visible at the protein level using immunohistochemistry in our experimental conditions. Consequently, it would be interesting to validate these results using more sensitive and comprehensive proteomic approaches.

The effect of a repeated LiFUS+MB treatment, 4 weekly sonication, was then investigated, as a chronic treatment protocol is more translationally relevant for AD. At an advanced stage of the disease, a short-term inflammatory response persisted up to 7 days after the last LiFUS+MB session. This response predominantly affected microglial cells at the transcriptional level, with modulation of *Aif1*, *P2ry12*, *Cx3cr1*, *Trem2*, and *C3* gene expression. To further assess long-term consequences, sorted microglia from aged rat brains were analyzed 6 weeks after the last sonication. At this advanced stage of the disease, LiFUS+MB did not induce major changes in microglia inflammatory profile and functions. The baseline level of microglia activation at this late pathological stage could be already elevated, potentially masking LiFUS-mediated effects. Interestingly, when the same repeated treatment was applied at an earlier disease stage, a modulation of microglia phenotype was observed with changes in the mRNA expression of several inflammatory markers, both pro- (e.g. *Tnfα*, *Tspo*) and anti-inflammatory (e.g. *Tgfβ*). Our data did not reflect a clear polarization toward either a classical pro- or anti-inflammatory phenotype. In recent years, the classification of activated microglia into these two distinct categories has become increasingly debated. Instead, it is now described as a continuum of transformation, with microglia continuously adapting to their environment and monitoring it through ongoing morphological and functional changes^52^. In our conditions, after a repeated LiFUS+MB treatment in younger animals, a decrease in the expression of *P2y12r*, a specific microglia marker that is expressed in their processes interacting with astrocytic endfeet, pericytes and a part of the endothelial cells in cortical blood vessels^53^, was observed. A modulation of the *P2y12r* expression is therefore consistent with our LiFUS+MB procedure to disrupt the BBB. Moreover, it has been shown that a low level of P2Y12R in microglia would be a marker of inflammatory around Aβ plaques in human brain^51,54^. Surprisingly, at both early and advanced stages of the disease, TSPO expression was decreased at the mRNA and protein levels 6 weeks after chronic LiFUS treatment, in contrast to the short-term effects observed after single or repeated LiFUS sonications. At the early pathological stage, this reduction of TSPO was accompanied by an increase in *Tnfα* mRNA levels, a downstream modulator of the NFκB pathway. This upregulation has already been observed in cultures of TSPO-deficient human microglia and a compensatory mechanism was proposed in favor of the NFκB pathway, which would additionally be negatively regulated by TSPO^55,56^. The discrepancy between short- and long-term effects of LiFUS exposure on TSPO expression would suggest that the early increase in *Tspo* levels was more likely associated with a transient inflammatory response, whereas the later reduction in TSPO may reflect a sustained functional adaptation of microglia than acute inflammatory activation.

Indeed, in addition to the inflammatory changes, modifications in the expression of genes involved in microglia key functions were observed after a repeated LiFUS treatment at early-stage pathology. A decreased expression of the mitochondrial gene, *Vdac1,* was notably measured. In recent years, a structural and functional interaction between VDAC1 and TSPO has been proposed and a reduction of the VDAC1 protein level is observed in TSPO-deficient human microglia culture which affected the intracellular Ca^2+^ homeostasis, important for the bioenergetic function of the cells^55,57^. Moreover, the slight changes in the inflammatory state of microglia after multiple LiFUS+MB exposure is also associated with a metabolic adaptation, with an upregulation of the glucose transporter *Glut1* gene expression, suggesting an increase of glucose uptake which may induce an upregulation of the glycolysis by microglia. Moreover, our result also showed a trend toward decreased *Mct1* mRNA level without significative changes in *Mct4* expression. MCT1 and MCT4 are both involved in the transport of lactate and ketone bodies but differ in substrate affinity and are preferentially implicated in lactate import for MCT1 and export for MCT4^58,59^. Our results would therefore suggest a metabolic shift in microglia after multiple exposure in favor of the glycolysis process compared to the anaerobic and ketolysis pathways, already described in inflammatory conditions^60–62^.

In addition to inflammation modulation, different studies in mouse models of AD described a reduction of amyloid plaques after LiFUS exposure alone^17–19,21–23,63–66^. The effects on tau are more controversial with studies describing a reduction, an increase or no effect on Tau levels after sonication, as reviewed by Géraudie *et al.*^67^. For the first time to our knowledge, the effect of LiFUS+MB was investigated on different poorly or highly aggregated forms of Aβ. Surprisingly, the impact of LiFUS+MB on the different forms of amyloid varied depending on delay post treatment, treatment schedule and disease stage when the treatment was applied. Indeed, in old animals, at short-term, LiFUS+MB only impacted Aβ42 whereas Aβ40 peptides are decreased at long term, suggesting a modulation of different aggregation or degradation pathways depending on the delay. Moreover, a chronic treatment even induced an increase of pTau231 levels in addition to aggregated Aβ42 forms at short-term, which appears to return to a normal long-term progression. The amyloid and Tau modulations are unaffected at early stage of the disease. Consequently, in our experimental conditions (LiFUS parameters, MB concentration, delay post sonication), by disrupting safely the BBB, LiFUS appears more promising as a key tool to improve drug delivery into the brain for AD, allowing the reduction of the i.v injected dose and reducing potentially the risk of side effects.

## Conclusion

Microglia are highly plastic cells capable of altering their phenotype and functions in response to environmental stimuli. Single or repeated LiFUS exposure induced a slight increase of glial reactivity at the transcriptional level at short-term in our experimental conditions. At early pathological stage, repeated LiFUS treatment seems to induce microglial reprogramming, leading to the adaptation of gene expression related to key functions such as inflammatory response, mitochondrial and energetic metabolism. Overall, LiFUS treatment did not induce major changes in AD pathological hallmarks but additional studies at other timepoints appear essential to better characterize their impact on neural cells and brain functioning in general.

## Acknowledgements

We thank Pia Lovero, Maria Surini, Adrien Fischer, Natacha Reich and Mayra Gaafar for their technical assistance. We are grateful for the contribution of the ‘Association IFRAD Suisse,’ created in 2009 at the initiative of the ‘Fondation pour la Recherche sur Alzheimer’ (formerly IFRAD France). We also are grateful to the ‘Société Académique de Genève’, the ‘Commission Administrative’ and the ‘Commission Informatique’ of Geneva University and the Ernest Boninchi foundation for their support in acquiring the LiFUS apparatus.

