## Supplemental figure 1 for "Repeated low-intensity focused ultrasound induces microglial profile changes in the TgF344-AD rat model of Alzheimer’s disease"

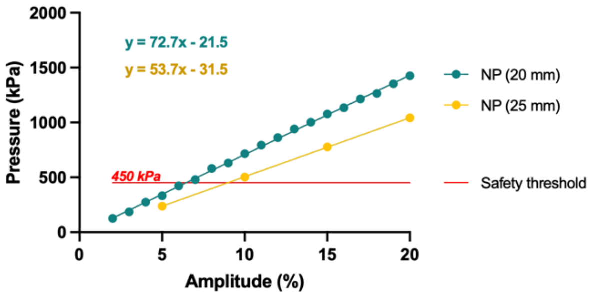


**Supplemental figure 1: Calibration of our LiFUS apparatus.**
To calibrate our apparatus for LiFUS+MB treatment, a submersible hydrophone was placed in degassed water below the transducer. The measurement of the peak negative acoustic pressure delivered by the transducer was realized at 20 (natural point) and 25 mm (steering). The maximal pressure used for all experiment was 450kPa.
