## Supplemental figure 2 for "Repeated low-intensity focused ultrasound induces microglial profile changes in the TgF344-AD rat model of Alzheimer’s disease"

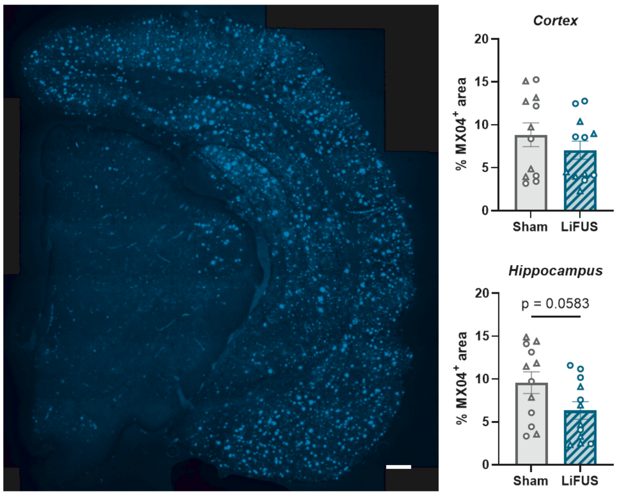


**Supplemental figure 2: Amyloid plaques are reduced 24h after LiFUS in the hippocampus.** Representative images of Methoxy-X04 (MX04; blue), labeling dense-core amyloid plaques. Scale bar = 500 µm. Quantification of the % $MX04^{+}$ area in the cortex and the hippocampus 24h after LIFUS+MB exposure. Cortex: Mann-Whitney test *p=0.4776;* Hippocampus: Mann-Whitney test *p=0.0583.* Males are represented by round points and females by triangles.
